# Mitochondria-derived dsRNA-induced stress granules promote IRF3-mediated fibrotic responses

**DOI:** 10.1101/2025.05.14.654103

**Authors:** Jared Travers, Emily Huang, Megan R. McMullen, Jianguo Wu, Christina K. Cajigas-Du Ross, Evi Paouri, Dimitrios Davalos, Vai Pathak, Daniel M. Rotroff, Laura E Nagy

## Abstract

Nucleic acid-induced activation of the anti-viral transcription factor interferon regulatory factor 3 (IRF3) promotes transforming growth factor-beta (TGFβ)-induced fibrotic responses of hepatic stellate cells *in vitro*. Herein, we identified molecular mechanisms underlying TGFβ-induced IRF3 activation. Gene silencing of *IRF3* with short interfering RNA (siRNA) in LX2 human hepatic stellate cells decreased TGFβ-induced fibrotic responses. An unbiased proteomic analysis of homeostatic IRF3-interacting proteins revealed striking enrichment in cytoplasmic stress granule components, including G3BP stress granule assembly factor 1 (G3BP1), G3BP2, and cell cycle associated protein 1 (CAPRIN1). TGFβ induced cytoplasmic accumulation of mitochondria-derived double-stranded RNA (mt-dsRNA) and assembly of IRF3-containing stress granules. Finally, blockade of G3BP1 activity or depletion of mt-dsRNA decreased TGFβ-induced fibrotic responses. Collectively, these findings identify a novel TGFβ-mt-dsRNA-stress granule axis in hepatic stellate cells that promotes IRF3-mediated fibrotic responses and potentially implicate cytoplasmic stress granules as a global regulator of fibrosis.

## INTRODUCTION

Fibrosis is a hallmark of advanced liver disease and involves excessive transdifferentiation and transactivation of quiescent hepatic stellate cells in response to a complex interplay of stimuli, including the multifunctional cytokine transforming growth factor-β (TGFβ)^1^. Liver fibrosis causes major morbidity and mortality and imposes a major financial burden on health care systems globally. Current treatment paradigms rely on orthotopic liver transplantation or preventing further fibrotic progression through a combination of abstinence from offending agents and risk factor modification^2^. The only FDA-approved pharmacologic agent for reversal of liver fibrosis, resmetirom, is only indicated in patients with Stage F2 or F3 fibrosis from metabolic dysfunction-associated steatotic liver disease^3^. Prevention and treatment of liver fibrosis is an unmet clinical need. Therefore, it is important to elucidate the factors that regulate hepatic stellate cell activation.

Mitochondria-derived endogenous nucleic acids are damage-associated molecular patterns (DAMPs) that trigger inflammatory responses. Mitochondria-derived DNA (mtDNA) is the most well-studied mitochondria-derived nucleic acid DAMP and is implicated in the pathogenesis of a multitude of human diseases^4–7^. Mitochondria-derived double-stranded RNA (mt-dsRNA) is continually created and degraded as part of the mtDNA replication process^4^. While less studied as a DAMP compared to mtDNA, mt-dsRNA has recently emerged as an important contributor to multiple disease states^8–10^. However, to the best of our knowledge, mt-dsRNA has not been linked to TGFβ-associated responses or fibrosis of any organ.

Interferon regulatory factor 3 (IRF3) is ubiquitously expressed and critically regulates host defense against viral infection^11,12^. IRF3 resides in the cytoplasm during homeostasis. Upon activation via multiple microbial byproducts or nucleic acid sensing pathways^13^, IRF3 translocates to the nucleus and induces transcription of type I interferons and other anti-viral genes^11,13^. In addition to this canonical transcription factor activity, IRF3 executes multiple non-transcriptional functions^14^. Activated IRF3 restrains Nuclear Factor kappa-B (NFκB) activity by direct binding to the p65 subunit in the cytoplasm^15^. Activated IRF3 also translocates to the mitochondrial outer membrane where it promotes apoptosis through direct interaction with Bax^16,17^. There is emerging evidence implicating both the transcriptional and non-transcriptional functions of IRF3 in the pathogenesis of metabolic liver diseases^12,18–20^, including alcohol-associated liver disease^21–23^. The contributions of IRF3 to the development of liver fibrosis is not clear as global *Irf3^-/-^* are reported to have both enhanced^24^ and diminished^25^ carbon tetrachloride (CCl_4_)-induced liver fibrosis. However, *in vitro* studies using isolated primary murine or immortalized human hepatic stellate cells suggest that hepatic stellate cell-intrinsic IRF3 activation mediates TGFβ-induced fibrogenic gene expression^26^ at least in part due to cytosolic sensing of mtDNA by the cGAS-STING pathway^26^. The potential contribution of other endogenous nucleic acids to IRF3-mediated fibrogenic responses in hepatic stellate cells has not, as yet, been assessed.

Cytoplasmic stress granules are large, organized, membrane-less conglomerates of untranslated mRNAs, stalled ribosomes, and RNA-binding proteins that form in response to various conditions of cellular stress^27^, including RNA virus infection^28^. Stress granules serve as a form of liquid-liquid phase separation to limit translation until the stressing event has passed, at which point they disassemble. Their assembly requires the actions of core nucleating proteins such as G3BP stress granule assembly factor 1 (G3BP1), its functional homolog G3BP2, and cell cycle associated protein 1 (CAPRIN1)^27^. Cytoplasmic stress granules are most well-studied in the context of anti-viral defense^28,29^. Emerging data has implicated stress granule formation in the pathogenesis of multiple cancers^30,31^ and neurodegenerative diseases^32–34^, but their potential involvement in fibrosis is unknown.

Herein, we investigated the profibrogenic activity of hepatic stellate cell-intrinsic IRF3 using the model fibrotic stimulus TGFβ. An unbiased proteomic analysis of IRF3-interacting proteins and three-dimensional (3D) immunofluorescence analysis revealed the presence of IRF3 inside of cytoplasmic stress granules induced by multiple stimuli. TGFβ induced cytoplasmic accumulation of mitochondria-derived dsRNA and assembly of IRF3-containing stress granules. Importantly, TGFβ-induced IRF3 activation and fibrogenic gene expression was diminished upon blockade of mitochondrial RNA synthesis or G3BP1 activity. These results identify TGFβ-induced mitochondrial dsRNA release and cytoplasmic stress granule assembly as important contributors to IRF3-mediated profibrotic responses of hepatic stellate cells.

## RESULTS

### IRF3 exerts pro-fibrotic effects *in vivo* and *in vitro*

Contradictory results in chronic carbon tetrachloride (CCl_4_)-induced fibrosis have been obtained with global *Irf3^-/-^* mice^24,25^. Therefore, we evaluated the contribution of *Irf3* to chronic CCl_4_-induced hepatic fibrosis. Significant hepatic fibrosis was induced in wild-type but to a lesser degree in *Irf3*^-/-^ mice as evidenced by Picrosirius Red staining in livers obtained 72 hours after the last CCl_4_ dose (Figure 1A,B). CCl_4_-induced expression of collagen 1 alpha 1 (*Col1a1*) mRNA (Figure 1C) and protein expression of the hepatic stellate cell activation marker α-smooth muscle actin (αSMA) (Figure 1D,E) were lower in *Irf3*^-/-^ mice as compared to WT. Collectively, these results reinforce an important role for IRF3 in chronic CCl_4_-induced hepatic fibrogenesis.

**Figure 1:**
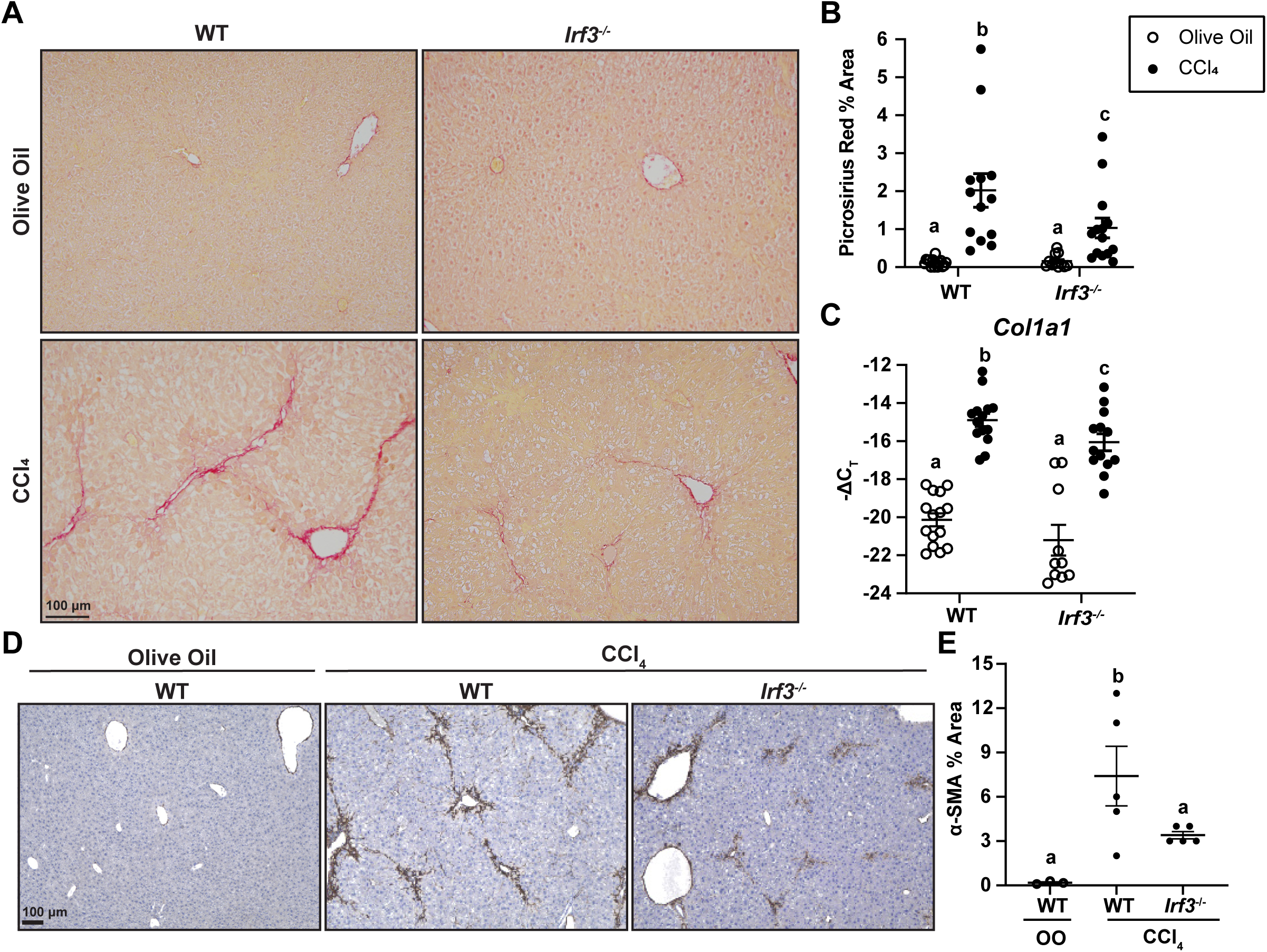
*Irf3*-deficient mice are protected against chronic CCl_4_-induced liver fibrosis. C57BL/6 and *Irf3^-/-^* mice were exposed to CCl_4_ or olive oil for five weeks with liver harvest 72 hours after the last dose. Picrosirius Red staining **(A,B)** or immunohistochemistry for α-SMA **(D,E)** were performed on paraffin-embedded liver sections. Images were obtained using a 10X objective, and the area of positive staining was quantified. **(C)** Expression of *Col1a1* mRNA in isolated livers was analyzed by qRT-PCR. C_T_ values were normalized to 18S rRNA and are shown as negative Δ C_T_ for a more intuitive data representation. For all graphs, each dot represents an individual mouse, and the error bars indicate the mean ± SEM. Values with different alphabetical superscripts are significantly different, P < 0.05.

In order to better understand the profibrogenic role of IRF3, *in vitro* responses of hepatic stellate cells to transforming growth factor-beta (TGFβ) were interrogated. Gene silencing of *IRF3* with short interfering RNA (siRNA) in LX2 human hepatic stellate cells decreased TGFβ-induced expression of fibrogenic genes, including *COL1A1* and α-smooth muscle actin (*ACTA2*) (Figure 2A), and accumulation of type I collagen in conditioned media (Figure 2B,C). 3D immunofluorescence analysis revealed that TGFβ stimulation modestly enhanced nuclear translocation of IRF3 (1.15-fold increase) (Figure 2D,E). Furthermore, gene silencing of the extracellular receptor for type I interferons (*IFNAR*) did not affect TGFβ-induced expression of fibrogenic genes (Figure 2F), indicating that the pro-fibrotic responses of LX2 cells to TGFβ do not depend on type I interferon production. Additionally, *IRF3* gene silencing did not affect gene expression of the two extracellular domains of the heterodimeric TGFβ receptor (*TGFBR1/2*) (Figure 2G).

**Figure 2:**
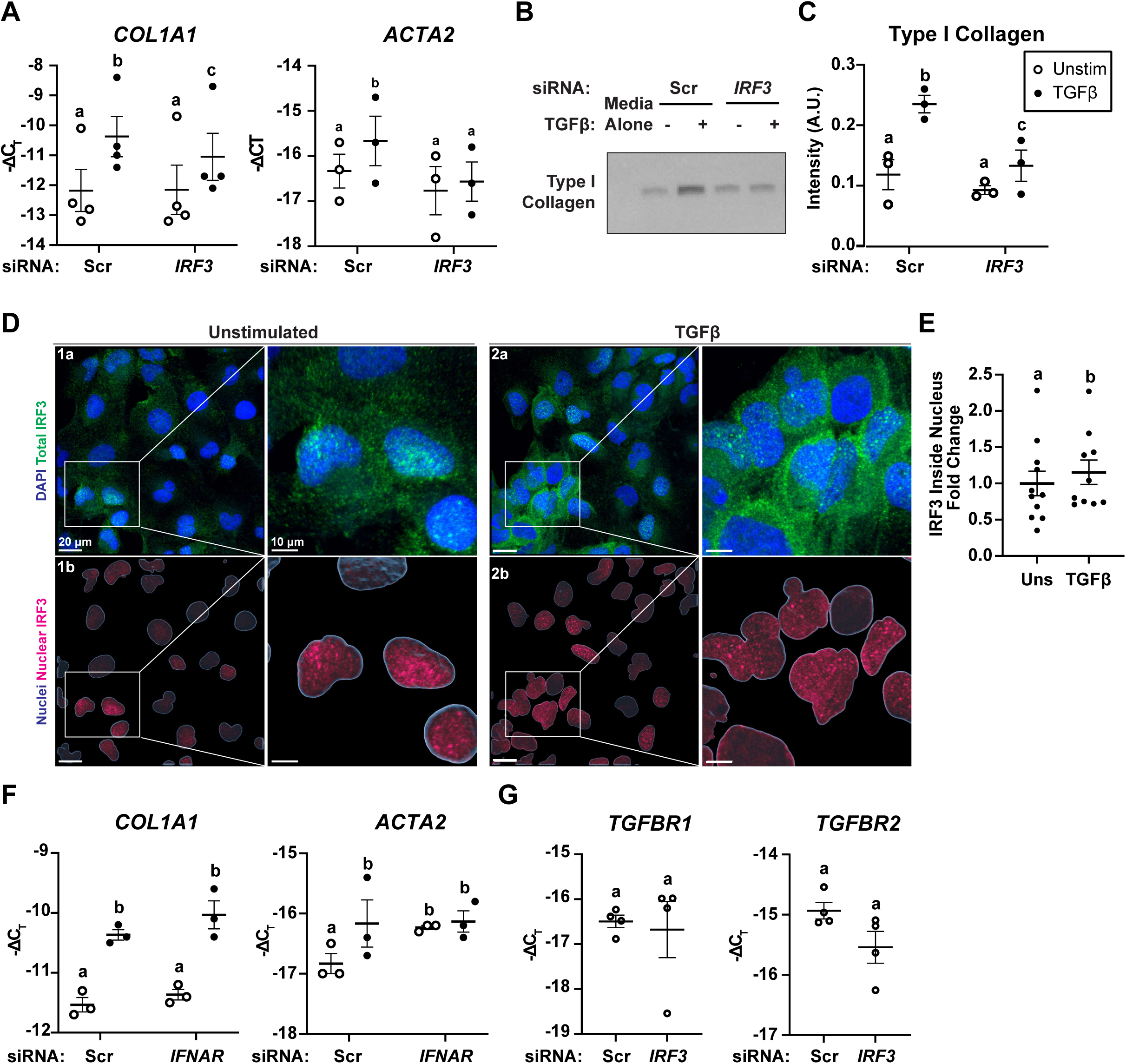
IRF3 promotes TGFβ-induced fibrotic responses in LX2 cells. **(A-C,F,G)** LX2 cells were transfected with scrambled control siRNA or siRNA against *IRF3* (**A-C,G**) or *IFNAR* (**F**) and then stimulated with 5 ng/mL TGFβ for 24 hours. **(A,F,G)** Expression of indicated mRNAs was assessed by qRT-PCR. C_T_ values were normalized to 18S rRNA and are shown as negative Δ C_T_ for a more intuitive data representation. **(B)** Western blot was performed on conditioned media to assess Type I collagen protein expression with quantification in (**C**). **(D-E)** LX2 cells were stimulated with 5 ng/mL TGFβ for 16 hours, then immunofluorescence was performed with anti-IRF3 antibody and DAPI to indicate nuclei. 3D reconstructions of Z-stacks were created with IMARIS, and the *Surfaces* module was used to depict nuclei. **(D)** Representative top-down views showing DAPI (blue, top row), total IRF3 (green, top row), nuclei (blue, bottom row), or IRF3 inside nuclei (pink, bottom row). Quantification of IRF3 fluorescence inside the nucleus is shown in (**E**). Data was normalized to the nuclear volume of the field and are depicted as fold change relative to unstimulated cells. For all graphs, each dot represents the mean of an individual experiment, and the error bars indicate the cumulative mean ± SEM. Values with different alphabetical superscripts are significantly different, P < 0.05.

### IRF3 is present within cytoplasmic stress granules in LX2 cells

Since type I interferon production was not required for TGFβ-induced fibrotic responses of LX2 cells, this prompted a search for novel regulators of IRF3’s profibrotic activity.

Immunoprecipitation was performed with an anti-IRF3 antibody in unstimulated LX2 cells, and the immunoprecipitates were characterized by mass spectrometry to find novel IRF3 binding partners. Over 500 immunoprecipitated proteins were identified (Supplementary Table 1), including a large number of DNA and RNA-binding proteins. The most abundant proteins included histones, polyadenylate-binding proteins, and the DNA polymerase delta catalytic subunit. Gene ontology enrichment analysis of immunoprecipitated proteins identified pathways involved in metabolism of RNA and cellular responses to stress (Figure 3A). Unexpectedly, several cytoplasmic stress granule component proteins were identified, including CAPRIN1, G3BP1, and G3BP2, eukaryotic translation initiation factor 2A (EIF2A), Fragile X messenger ribonucleoprotein 1 (FMR1), Fragile X-related protein 1 (FXR1), ELAV like RNA binding protein 1 (ELAVL1), and DEAH-box helicase 36 (DHX36) (Figure 3A). Despite robust detection in the mass spectrometry assay, in co-immunoprecipitation experiments, a direct physical association between IRF3 and the core stress granule protein G3BP1 was not detectable by Western blot at baseline (Figure 3B) or after transfection with the dsRNA analogue poly(I:C) (Figure 3C), a prototypical inducer of both IRF3 activation^13^ and stress granule assembly^35^.

**Figure 3:**
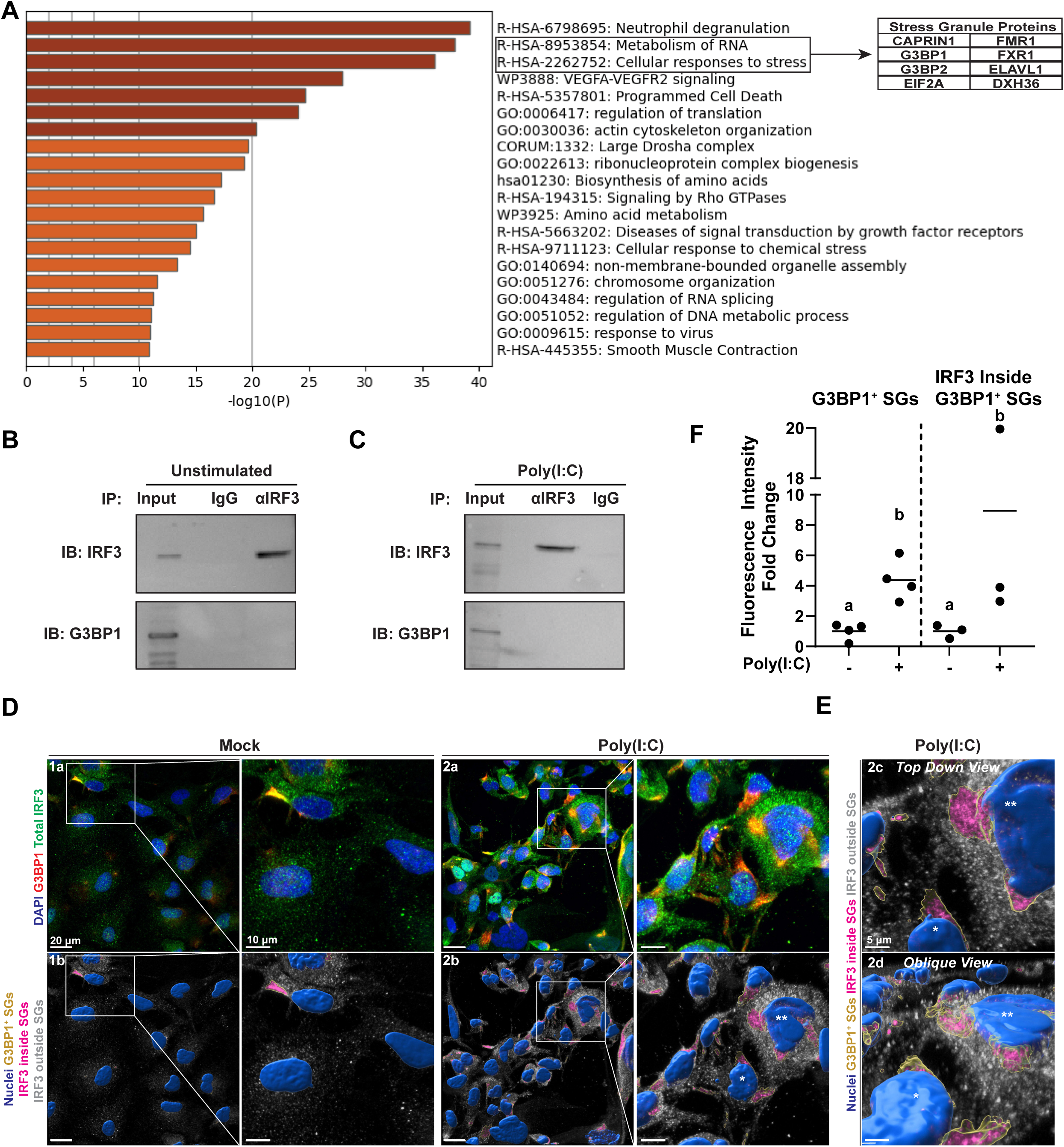
IRF3 is present within poly(I:C)-induced stress granules in LX2 cells. **(A)** Immunoprecipitation was performed with an anti-IRF3 antibody in unstimulated LX2 cells, and the immunoprecipitates were subjected to mass spectrometry. **(A)** Gene Ontology enrichment analysis of immunoprecipitated proteins. Dark orange bars indicate P < 10^-20^, and light orange bears indicate P < 10^-10^. Table lists example SG component proteins that were immunoprecipitated. **(B,C) I**mmunoprecipitation was performed with anti-IRF3 antibody or isotype control followed by Western blotting for IRF3 and G3BP1 in LX2 cells that were either unstimulated (**B**) or transfected with 1 μg/mL of poly(I:C) for four hours (**C**). Data are representative of three independent experiments. **(D-F)** LX2 cells were mock-transfected or transfected with 1 μg/mL poly(I:C) for 4 hours, then immunofluorescence was performed with anti-IRF3 antibody, anti-G3BP1 antibody, and DAPI to indicate nuclei. 3D reconstructions of Z-stacks were created with IMARIS, and the *Surfaces* module was used to depict nuclei and G3BP1^+^ SGs. **(D)** Subpanels 1a and 2a are representative top-down views showing DAPI (blue), G3BP1 (red), and total IRF3 (green), and subpanels 1b and 2b show nuclei (blue), G3BP1^+^ SGs (yellow), and IRF3 inside (magenta) and outside (gray) of G3BP1^+^ SGs. **(E)** Higher-magnification top-down and oblique views of inset from subpanel 2b. For orientation purposes, certain nuclei are labeled with asterisks. **(F)** Quantification of G3BP1 aggregation and IRF3 accumulation inside G3BP1^+^ SGs. Data was normalized to the nuclear volume of the field and are depicted as fold change relative to unstimulated cells. Each dot represents the mean of an individual experiment, and the horizontal line indicates the cumulative mean. Values with different alphabetical superscripts are significantly different, P < 0.05.

Immunofluorescence followed by confocal microscopy and 3D image analysis was then performed to determine the subcellular localization of IRF3 and G3BP1. Focal cytoplasmic G3BP1 aggregates, which are indicative of stress granule assembly^27^, were occasionally observed with mock transfection but markedly increased with poly(I:C) transfection (Figure 3D subpanels 1a/2a,F). Poly(I:C) transfection also induced punctate cytoplasmic aggregation of IRF3 (Figure 3D subpanels 1a/2a), which were confirmed to be inside G3BP1+ stress granules (Figure 3D subpanels 1b/ 2b,F) using 3D reconstructions and masking of the IRF3 fluorescence channel with IMARIS software. Furthermore, high-magnification top-down and oblique images illustrated that IRF3 was not distributed uniformly within G3BP1^+^ stress granules, but present as punctate foci with higher IRF3 fluorescence (Figure 3E subpanels 2c/2d). These findings identify the presence of IRF3 within cytoplasmic stress granules in response to poly(I:C), a classical IRF3 activation signal.

### TGFβ induces assembly of IRF3-containing stress granules in LX2 cells

Because IRF3 is profibrotic and localizes to poly(I:C)-induced cytoplasmic stress granules, we used 3D immunofluorescence analysis to test the hypothesis that IRF3-containing stress granules are also formed in response to TGFβ. Occasional cytoplasmic G3BP1^+^ or CAPRIN1^+^ stress granules were identified at baseline (Figure 4A/C subpanels 1a/1b). TGFβ strongly induced cytoplasmic aggregation of both G3BP1 and CAPRIN1 (Figure 4A/C subpanels 2a/2b, E,F). IRF3 was detectable throughout G3BP1^+^ and CAPRIN1^+^ stress granules (Figure 4A/C subpanels 1b/2b,E,F). Punctate areas of higher IRF3 aggregation were visible both inside and immediately surrounding stress granules in high-magnification views (Figure 4B,D). Taken together, these findings identify TGFβ as a novel inducer of the assembly of IRF3-containing stress granules.

**Figure 4:**
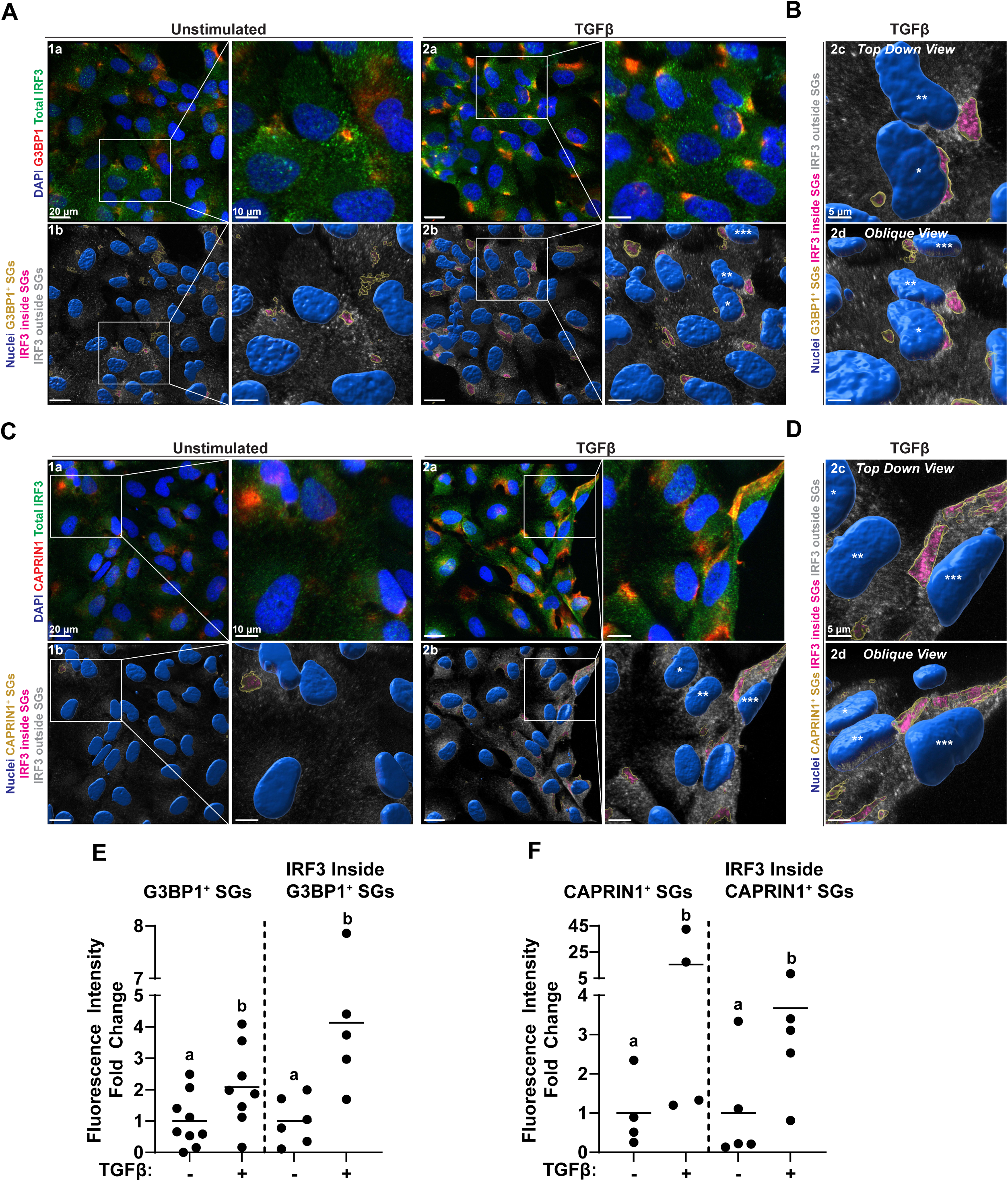
TGFβ induces assembly of IRF3-containing stress granules in LX2 cells. LX2 cells were stimulated with 5 ng/mL of TGFβ for 16 hours, then immunofluorescence was performed with anti-IRF3 antibody, anti-G3BP1 (**A,B**) or anti-CAPRIN1 (**C,D**) antibody, and DAPI to indicate nuclei. 3D reconstructions of Z-stacks were created with IMARIS, and the *Surfaces* module was used to depict nuclei and G3BP1^+^ or CAPRIN1^+^ SGs. **(A,C)** Subpanels 1a and 2a are representative top-down views showing DAPI (blue), G3BP1 or CAPRIN1 (red), and total IRF3 (green), and subpanels 1b and 2b show nuclei (blue), G3BP1^+^ or CAPRIN1^+^ SGs (yellow), and IRF3 inside (magenta) and outside (gray) of G3BP1^+^ or CAPRIN1^+^ SGs. **(B,D)** Higher-magnification top-down and oblique views of insets from subpanels **A**2b and **C**2b. For orientation purposes, certain nuclei are labeled with asterisks. **(E,F)** Quantifications of G3BP1 (**E**) or CAPRIN1 (**F**) aggregation and IRF3 accumulation inside G3BP1^+^ (**E**) or CAPRIN1^+^ (**F**) SGs. Data was normalized to the nuclear volume of the field and are depicted as fold change relative to unstimulated cells. For all graphs, each dot represents the mean of an individual experiment, and the horizontal line indicates the cumulative mean. Values with different alphabetical superscripts are significantly different, P < 0.05.

### Cytoplasmic stress granules promote fibrotic responses in LX2 cells

We next determined the impact of stress granules on fibrotic responses to TGFβ using both pharmacologic and genetic approaches. Pharmacologic inhibition of G3BP1 activity with (-)- epigallocatechin gallate (EGCG)^36–38^ decreased expression of the fibrogenic genes *COL1A1* and *FN1* and accumulation of type I collagen and fibronectin in conditioned media both at baseline and upon TGFβ stimulation (Figure 5A-C). Genetic blockade of G3BP1 was also performed to corroborate these findings. LX2 cells were co-transfected with siRNA targeting *G3BP1* and its functional homolog *G3BP2*. Efficient gene silencing of both *G3BP1* and *G3BP2* was confirmed by Western blotting (Supplementary Figure 1). *G3BP1*/*G3BP2* dual gene-silencing also decreased TGFβ-induced *COL1A1* and *FN1* mRNA expression and accumulation of type I collagen and fibronectin in conditioned media (Figure 5D-F).

**Figure 5:**
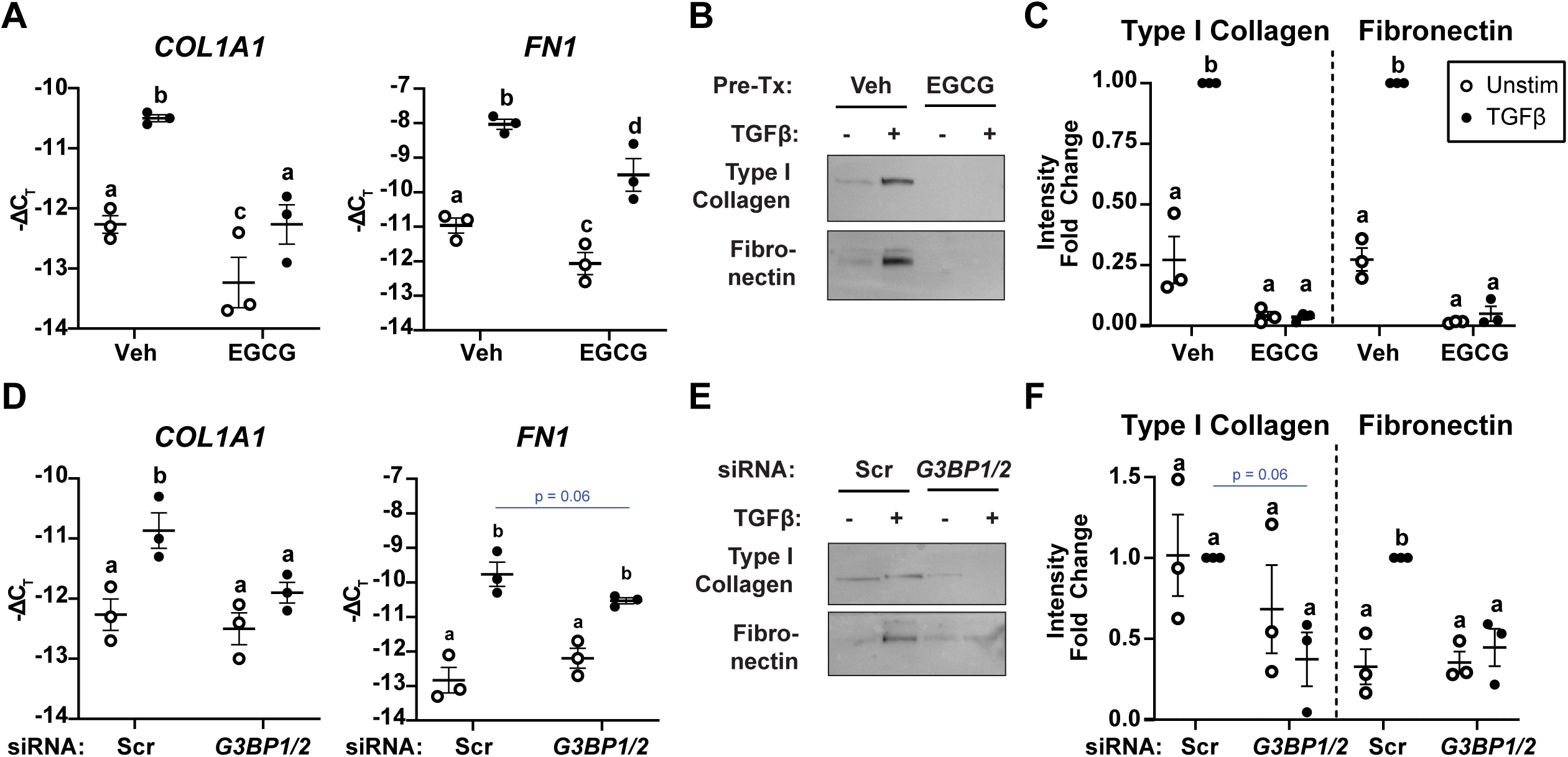
Cytoplasmic stress granules promote fibrotic responses to TGFβ in LX2 cells. LX2 cells were (**A-C**) pre-treated for one hour with 25 μM EGCG or vehicle or (**D-F**) transfected with scrambled control siRNA or siRNAs against *G3BP1* and *G3BP2* and then stimulated with 5 ng/mL TGFβ for 24 hours. **(A,D)** Expression of *COL1A1* and *FN1* mRNA was analyzed by qRT-PCR. Values were normalized to 18S rRNA and are shown as negative Δ C_T_ for a more intuitive data representation. (**B,E)** Western blot was performed on conditioned media to assess Type I collagen and fibronectin protein expression with quantification in (**C**) and (**F**), respectively. For all graphs, each dot represents the mean of an individual experiment and the error bars indicate the cumulative mean ± SEM. Values with different alphabetical superscripts are significantly different, P < 0.05.

### TGFβ induces accumulation of mitochondrial dsRNA in LX2 cells

The mechanism underlying TGFβ-induced stress granule assembly was then investigated. Intracellular foreign nucleic acids, such as viral dsRNA, are well-described inducers of stress granule assembly^27,35^ and IRF3 activation^13^. However, the functional links between endogenous dsRNA accumulation and IRF3 activation are not well understood. Therefore, the hypothesis that TGFβ stimulation results in accumulation of dsRNA to activate IRF3 was examined. 3D immunofluorescence analysis utilizing a sequence-agnostic dsRNA monoclonal antibody (J2) revealed intracellular dsRNA accumulation upon TGFβ treatment (Figure 6A subpanels 1a/2a,E). TGFβ increased dsRNA inside of G3BP1^+^ stress granules (Figure 6A subpanels 1b/2b,E). 3D co-localization revealed that a significant proportion of dsRNA (approximately 8%) was present inside of G3BP1+ stress granules. Higher magnification views revealed that dsRNA is not present throughout the entire stress granule but rather only in specific foci (Figure 6B). Because nuclei and mitochondria are by far the most common origin of endogenous dsRNA^39^, their co-localization with accumulated dsRNA was assessed with DAPI staining and fluorescent mitochondrial labeling (MitoTracker), respectively (Figure 6C,D). While TGFβ increased focal accumulation of both mitochondrial and nuclear dsRNA (Figure 6E), only a small proportion of dsRNA was present inside the mitochondria and nucleus (approximately 11% and 23%, respectively) (Figure 6F). These results indicate that the accumulation of dsRNA was not related to retention within mitochondria or nuclei.

**Figure 6:**
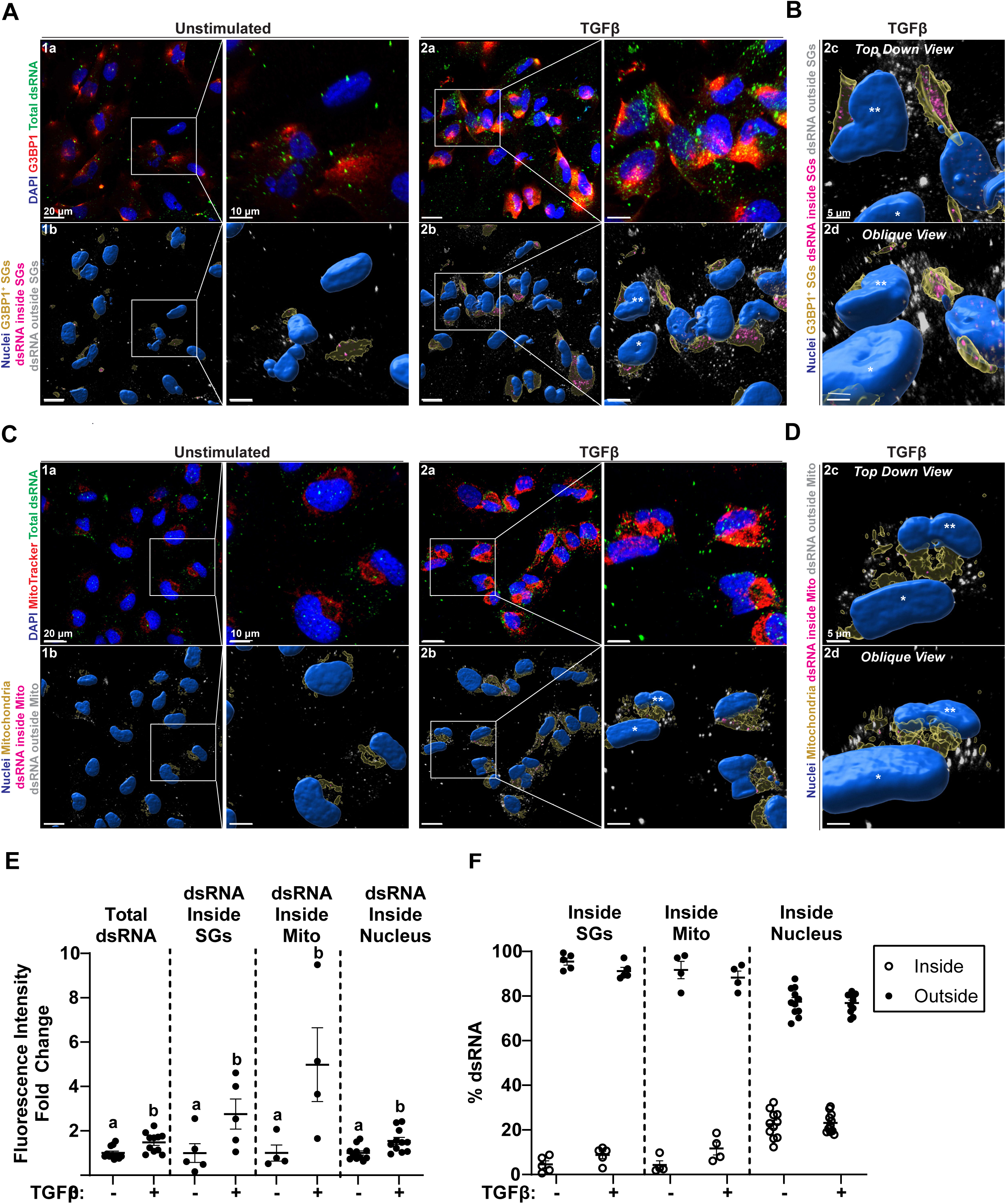
TGFβ induces accumulation of mitochondria-derived dsRNA in LX2 cells. LX2 cells were stimulated with 5 ng/mL of TGFβ for 16 hours, then immunofluorescence was performed with anti-dsRNA antibody (J2), anti-G3BP1 (**A,B**) or MitoTracker (**C,D**), and DAPI to indicate nuclei. 3D reconstructions of Z-stacks were created with IMARIS, and the *Surfaces* module was used to depict nuclei, mitochondria, and G3BP1^+^ SGs. **(A,C)** Subpanels 1a and 2a are representative top-down views showing DAPI (blue), G3BP1 or MitoTracker (red), and total dsRNA (green), and subpanels 1b and 2b show nuclei (blue), G3BP1^+^ SGs or mitochondria (yellow), and dsRNA inside (magenta) and outside (gray) of G3BP1^+^ SGs or mitochondria. **(B,D)** Higher-magnification top-down and oblique views of insets from subpanels **A**2b and **C**2b. For orientation purposes, certain nuclei are labeled with asterisks. **(E)** Quantifications of the total amount of dsRNA, dsRNA inside SGs, dsRNA inside mitochondria, and dsRNA inside nuclei. Data was normalized to the nuclear volume of the field and are depicted as fold change relative to unstimulated cells. **(F)** Quantification of the percentage of dsRNA inside or outside of G3BP1^+^ SGs, mitochondria, or nuclei. For all graphs, each dot represents the mean of an individual experiment, and the horizontal line indicates the cumulative mean ± SEM. Values with different alphabetical superscripts are significantly different, P < 0.05.

### Mitochondrial dsRNA mediates TGFβ-induced stress granule assembly and fibrotic responses

The functional impact of TGFβ-induced accumulation of mitochondrial dsRNA was examined next. Depletion of mitochondrial dsRNA with the mitochondrial RNA polymerase pharmacologic inhibitor 2’-C-methyladenosine (2-CM)^4^ decreased TGFβ-stimulated dsRNA accumulation (Figure 7A-C). Similarly, 2-CM pre-treatment also decreased TGFβ-induced G3BP1^+^ stress granule formation and the amount of dsRNA inside of stress granules (Figure 7A-C). 2-CM exerted no effect on dsRNA accumulation or G3BP1 aggregation in the absence of TGFβ stimulation. Depletion of mitochondrial dsRNA with 2-CM decreased TGFβ-induced expression of multiple fibrogenic genes, including *COL1A1*, *ACTA2*, and *FN1* (Figure 7D-F), and accumulation of the extracellular matrix proteins type I collagen and fibronectin in conditioned media (Figure 7G,H). As a whole, these findings show that TGFβ-induced release of mitochondrial dsRNA promotes stress granule formation and fibrotic responses.

**Figure 7:**
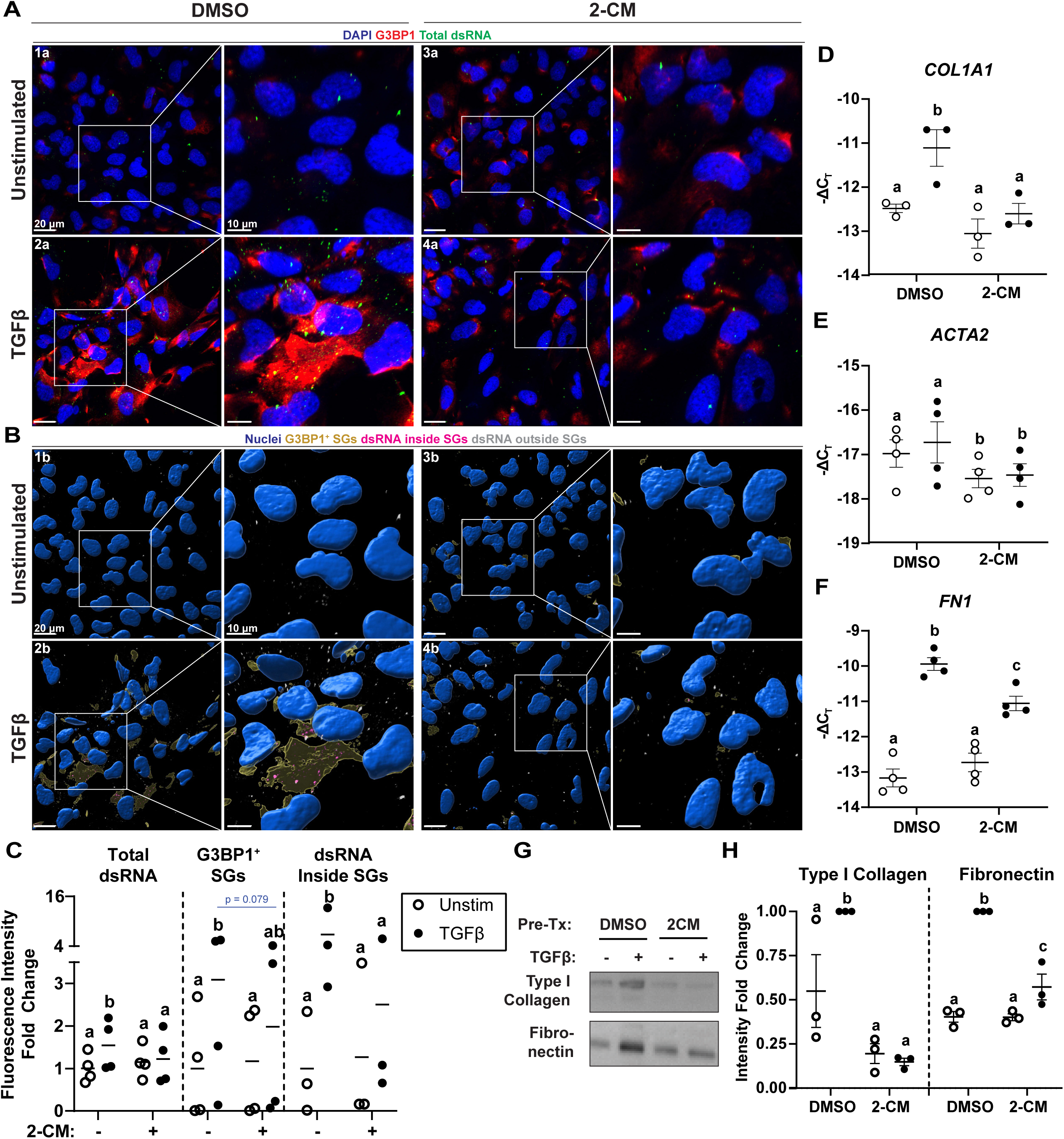
Depletion of mitochondria-derived dsRNA inhibits TGFβ-induced stress granule assembly and fibrotic responses in LX2 cells. LX2 cells were pre-treated for 24 hours with 20 μM of 2-CM or DMSO, then stimulated with 5 ng/mL of TGFβ for 16 hours (**A-C**) or 24 hours (**D-H**). **(A-C)** Immunofluorescence was performed with anti-dsRNA antibody (J2), anti-G3BP1 antibody, and DAPI to indicate nuclei. 3D reconstructions of Z-stacks were created with IMARIS, and the *Surfaces* module was used to depict nuclei, mitochondria, and G3BP1^+^. **(A,B)** Representative top-down views showing DAPI (**A,** blue), G3BP1 (**A,** red), total dsRNA (**A,** green), nuclei (**B,** blue), G3BP1^+^ SGs (**B,** yellow), and dsRNA inside (**B,** magenta) and outside (**B,** gray) of G3BP1^+^ SGs. **(C)** Quantifications of the total amount of dsRNA, total amount of G3BP1^+^ SGs formed, and dsRNA inside of G3BP1^+^ SGs. Data was normalized to the nuclear volume of the field and are depicted as fold change relative to DMSO-pretreated, unstimulated cells. **(D-F)** Expression of indicated mRNAs was analyzed by qRT-PCR. Values were normalized to 18S rRNA and are shown as negative Δ C_T_ for a more intuitive data representation. **(G)** Western blot was performed on conditioned media to assess Type I collagen and fibronectin protein expression with quantification in (**H**). For all graphs, each dot represents the mean of an individual experiment. The error bars indicate the cumulative mean ± SEM [except (**C**)]. Values with different alphabetical superscripts are significantly different, P < 0.05.

## DISCUSSION

IRF3, a key regulator of anti-viral defense, is increasingly appreciated to be important in metabolic liver diseases. However, the role of IRF3 specifically in liver fibrosis is unclear. While IRF3 promotes fibrotic responses of cultured hepatic stellate cells to TGFβ^26^, contradictory results in chronic CCl_4_-induced liver fibrosis are reported using global *Irf3^-/-^* mice^24,25^. The *in vivo* results of our study reinforce a profibrogenic role of IRF3. Moreover, our investigation into IRF3-mediated responses of cultured hepatic stellate cells to TGFβ led to multiple novel observations regarding the biology of cytoplasmic stress granules and mt-dsRNA. We discovered that TGFβ induces cytoplasmic accumulation of mt-dsRNA, which triggers assembly of IRF3-containing stress granules. Both mt-dsRNA and cytoplasmic stress granules were required for optimal TGFβ-induced fibrogenic gene expression. These findings illustrate novel pro-fibrotic roles for mt-dsRNA and IRF3-containing stress granules.

Our results complement both *in vivo*^24,25^ and *in vitro*^26^ lines of evidence implicating IRF3 in liver fibrosis. However, the mechanisms by which IRF3 exerts these effects remains poorly understood. Classically, upon activation IRF3 is phosphorylated, dimerizes, and then translocates to the nucleus where it enhances transcription of anti-viral genes like type I interferons and interferon-stimulated genes^13^. Indeed, while TGFβ stimulation was reported to increase IRF3 phosphorylation in LX2 cells^26^, we observed very modest IRF3 nuclear translocation in response to TGFβ stimulation in our study. Furthermore, gene silencing against the type I interferon receptor IFNAR did not affect TGFβ-induced fibrogenic gene expression. These results are consistent with a potential non-transcriptional mechanism of action for IRF3. Of note, IRF3 has been shown to directly interact with the TGFβ signaling molecule mothers against decapentaplegic homolog 3 (SMAD3)^40^, but this interaction has not as yet been assessed in hepatic stellate cells. Further investigation is needed to clarify the relative contributions of IRF3 transcriptional and non-transcriptional activities to fibrotic responses in hepatic stellate cells and liver fibrosis.

Our discovery that IRF3 is present within dsRNA-induced stress granules is supported by both proteomic and immunofluorescence-based subcellular localization analyses. Our mass spectrometry-based proteomic analysis of IRF3-interacting proteins adds stress granule component proteins to the list of known extranuclear IRF3 binding partners like Bax^16,17^, the p65 subunit of NFκB^15^, and SMAD3^40^. The results of our proteomic analysis were independently corroborated using three-dimensional immunofluorescence imaging. The presence of IRF3 within stress granules, rather than focal non-SG aggregates like G3BP1-cGAS complexes^37,38^, is supported by the findings that (1) IRF3 was localized to focal aggregates of multiple bonafide stress granule proteins (G3BP1 and CAPRIN1) and (2) TGFβ-induced G3BP1 aggregates contained dsRNA. Notably, IRF3 was not uniformly distributed throughout G3BP1^+^ and CAPRIN1^+^ SGs, and punctate areas of higher IRF3 aggregation were also detectable in the immediately surrounding areas. The main structural features of stress granules are dense cores, where ribonucleoproteins are tightly bound to nucleating proteins like G3BP1 and CAPRIN1^27^, and outer shells where weaker, more transient protein-protein interactions occur^27^. IRF3 may be predominantly present within the stress granule shell, rather than the core. This is a potential explanation for why a direct interaction between IRF3 and G3BP1 was not detectable in independent co-immunoprecipitation experiments by Western blot, which has a significantly higher detection limit than mass spectrometry.

Both pharmacologic and genetic blockade strategies were employed in our study to demonstrate that stress granules promote *in vitro* fibrotic responses to TGFβ. The pharmacologic G3BP1 activity inhibitor, EGCG, is a component of green tea extract^41^ and exerted anti-fibrogenic effects in multiple experimental rodent models of hepatic fibrosis^42–44^. The *in vitro* effects of EGCG were phenocopied by dual gene silencing of both *G3BP1* and its functional homolog *G3BP2*, which rules against off-target effects of the inhibitor. The mechanisms by which stress granules regulate the fibrotic response to TGFβ are unknown. While stress granules are most well-known for regulating mRNA translation by sequestering mRNA and translation initiation components^27^, they also serve as a platform for immune and stress signaling. Stress granules have been demonstrated to contain multiple nucleic acid sensors, such as retinoic acid-inducible gene I (RIG-I)^45,46^, melanoma differentiation-associated protein 5 (MDA5)^45,47^, and protein kinase R (PKR)^45,48^. Furthermore, stress granules can sequester intracellular signaling molecules, such as c-Jun N-terminal kinase (JNK)^28,49^ and mammalian target of rapamycin complex 1 (mTORC1)^28,50,51^, and nucleocytoplasmic transport proteins like importins^52–54^. Whether stress granules promote pro-fibrotic responses to TGFβ by dysregulating mRNA translation, enhancing the sensing of cytosolic endogenous nucleic acids, or directly modifying relevant signaling pathways remains unclear.

An important novel finding of our study is that TGFβ induces intracellular accumulation of dsRNA. Immunofluorescence-based assessment of the subcellular localization of accumulated dsRNA was consistent with cytoplasmic accumulation, rather than merely nuclear or mitochondrial retention. A predominantly mitochondrial origin of the accumulated dsRNA was identified through use of a pharmacologic inhibitor of the mitochondrial RNA polymerase to deplete mitochondria-derived dsRNA. We also examined the functional importance of the accumulated dsRNA. Depletion of mitochondria-derived dsRNA impaired both stress granule assembly and the fibrotic response to TGFβ. These results suggest that mitochondrial, rather than nuclear, dsRNA is more pro-fibrotic. Our finding of TGFβ-induced cytoplasmic accumulation of mitochondria-derived dsRNA parallels similar findings for mtDNA, which activated the cGAS-STING-IRF3 axis to promote stellate cell activation *in vitro*^26^. Further investigation is needed to elucidate the required proximal mt-dsRNA pattern recognition receptors and the events that occur upstream of mitochondrial nucleic acid release.

Both stress granules and mt-dsRNA are increasingly appreciated as important contributors to diverse physiological and pathological processes. Stress granules are most well-studied in the context of host defense against viral infection^28,29^ but also contribute to the pathogenesis of cancer^30,31^ and neurodegenerative diseases^33,34^. Although much less well-studied as a DAMP than mtDNA, mt-dsRNA is implicated in multiple inflammatory disorders^8,9^ as well as in metabolic regulation and differentiation of beige adipocytes^55^. Additionally, recent reports have separately implicated both stress granules and mt-dsRNA in cellular senescence^56,57^. To the best of our knowledge, this is the first report of a pro-fibrotic role for stress granules or mt-dsRNA.

In total, our investigation into the pro-fibrotic activity of IRF3 led to the discovery that TGFβ induces cytoplasmic accumulation of mitochondria-derived dsRNA, which triggers assembly of IRF3-containing stress granules to promote fibrotic responses. Given the ubiquitous nature of stress granules and mt-dsRNA, we expect our findings to spur further investigation into their contributions to other fibrotic and non-fibrotic disease states both in the liver and possibly in other organs as well.

## METHODS

### Mouse handling and tissue collection

All procedures using animals were approved by the Cleveland Clinic Institutional Animal Care and Use committee. Male and female C57BL/6 mice were purchased from Jackson Laboratory (Bar Harbor, ME). *Irf3*^-/-^ were obtained from Ganes Sen (Cleveland Clinic)^16^. 10-12 week-old male and female C57Bl/6 WT and *Irf3^-/-^* mice received intraperitoneal injections with CCl_4_ or olive oil as the vehicle control twice-weekly for five weeks. CCl_4_ was diluted 1:3 in olive oil. The initial dose was 0.25 μL/g body weight, the second was 0.5 μL/g body weight, followed by 6-8 additional doses at the full 1 μL/g body weight. Liver harvest occurred 72 hours after the final CCl_4_ exposure. Portions of liver were flash-frozen in liquid nitrogen and stored at −80 °Celsius (°C) or fixed in 10% formalin for histology.

### Picrosirius red staining and α-Smooth Muscle Actin immunohistochemistry

Formalin-fixed tissues were paraffin-embedded and sectioned with a thickness of 5 μm. After de-paraffinization, sections underwent Picrosirius red staining (365548, Sigma-Aldrich, St. Louis, MO) for histological detection of fibrosis or 3,3’-diaminobenzidine (DAB) chromogenic immunohistochemistry to detect α-SMA (1:200, 19245, Cell Signaling Technologies, Danvers, MA) protein expression. Slides were coded prior to initial examination. Quantification of picrosirius red staining and α-SMA IHC was performed using ImageJ software (NIH, Bethesda, MD). Images were taken at 10x and are representative of 3-5 mice per group. Values represent percent area stained for picrosirius red or α-SMA.

### Cell culture

LX2 immortalized human hepatic stellate cells were a gift from Scott Friedman (Mount Sinai, New York, USA)^58^ and were cultured in Dulbecco’s modified Eagle’s medium (11965092, DMEM; Thermo Fisher Scientific, Waltham, MA) supplemented with 2% fetal bovine serum (16000044, Thermo Fisher Scientific, Waltham, MA) and 1% L-glutamine (25030081, Thermo Fisher Scientific, Waltham, MA). This same media was used for all stimulations. For poly(I:C) stimulations, cells were transfected with Lipofectamine RNAiMax transfection reagent (13778150, Thermo Fisher Scientific, Waltham, MA) ± 1 μg/mL poly(I:C) (P1530, Sigma, St. Louis, MO) following the manufacturer instructions. For TGFβ stimulations, cells were given fresh culture media ± 5 ng/mL TGFβ1 (7666-MB-005, R&D, Minneapolis, MN) for indicated lengths of time. For pharmacologic inhibitor experiments, cells were pre-treated with 25 μM EGCG (4143, Sigma, St. Louis, MO) for one hour or 50 μM 2-CM (sc-283467, Santa Cruz, Dallas, TX) or DMSO vehicle control (N182, VWR, Radnor, PA) for 24 hours prior to TGFβ stimulation.

### Short interfering RNA transfection

After plating in 6-well or 12-well plates, cells were transfected the same day with scrambled control siRNA (sc-37007, Santa Cruz, Dallas, TX) or a pool of three siRNAs against human *IRF3* (encodes IRF3 protein) (SR320690, Origene, Rockville, MD) or human *IFNAR* (encodes Type I interferon receptor) (sc-35637, Santa Cruz, Dallas, TX) using Lipofectamine RNAiMAX transfection reagent following the manufacturer’s instructions. TGFβ stimulation was then performed 48 hours later. In other experiments, cells were transfected with scrambled control siRNA or a pool of three siRNAs against human *G3BP1* (encodes G3BP1 protein) (sc-75076, Santa Cruz, Dallas, TX). The next day, cells were transfected with scambled control siRNA or a pool of three siRNAs against human *G3BP2* (encodes G3BP2 protein) (sc-89231, Santa Cruz, Dallas, TX). TGFβ stimulation was then performed 24 hours later.

### RNA isolation and quantitative real-time polymerase chain reaction

Total RNA was isolated from frozen mouse liver specimens using RNeasy Plus Universal Mini Kit (73404, Qiagen, Germantown, MD) following the manufacturer’s instructions. Total RNA from cultured LX2 cells was isolated using Direct-zol RNA MicroPrep kit (R2062, Zymo Research, Irvine, CA) following the manufacturer’s instructions. After RNA isolation, complementary DNA (cDNA) was then synthesized using Superscript IV VILO Master Mix (11756500, Thermo Fisher Scientific, Waltham, MA). Quantitative RT-PCR was then performed in a Quantstudio5 real-time PCR system (Thermo Fisher Scientific, Waltham, MA) using Power SYBR PCR Master Mix (4367659, Applied Biosystems, Waltham, MA) and gene-specific primers (Supplementary Table 2). The relative amount of target mRNA was determined using the comparative threshold (C_T_) method by normalizing target mRNA Ct values to those of *18S*. Statistical tests were performed using ΔC_T_ values (average C_T_ of gene of interest minus average C_T_ of *18S*).

### Antibodies

Rabbit anti-fibronectin (26836T), rabbit anti-β-actin (4967S), and mouse IgG1 isotype control (5415) were purchased from Cell Signaling Technologies (Danvers, MA). Goat anti-Type I Collagen (1310-01) was purchased from Southern Biotech (Birmingham, AL). Mouse anti-IRF3 (sc-33641) were purchased from Santa Cruz (Dallas, TX). Rabbit anti-G3BP1 (13057-2-AP), rabbit anti-G3BP2 (16276-1-AP), and rabbit anti-CAPRIN1 (15112-1-AP) were purchased from Proteintech (Rosemont, IL). Mouse anti-GAPDH (MAB374) was purchased from MilliporeSigma (St. Louis, MO). Mouse anti-dsRNA [J2] (10010200) antibody was purchased from Scicons (Budapest, Hungary). HRP-labeled donkey anti-rabbit (A16035), goat anti-mouse (G21040), AlexaFluor 488-labeled goat anti-mouse (A11029), and AlexaFluor 568-labeled goat anti-rabbit (A11036) secondary antibodies were purchased from Invitrogen (Waltham, MA). HRP-labeled mouse anti-rabbit IgG (light chain-specific) (211-032-171) and goat anti-mouse IgG (light chain-specific (115-035-174) were purchased from Jackson ImmunoResearch (West Grove, PA).

### Western blot analysis

After washing with PBS, cultured LX2 cells were lysed in RIPA buffer (50 mM Tris-Hcl pH 8.0, 150 mM NaCl, 1% NP-40, 0.5% sodium deoxycholate, 0.1% SDS, 1 mM EGTA) supplemented with Protease and Phosphate Inhibitors (A32959, Thermo Fisher Scientific, Waltham, MA).

Protein concentrations were measured using DC Protein Assay (5000111, BioRad, Hercules, CA), then equal amounts of protein was resuspended in Laemmli Buffer, boiled at 95 °C for 5 minutes, and then resolved on either an 8% or 10% SDS-PAGE gel. After semi-dry transfer, membranes were blocked with 3% BSA in TBS-T then probed with primary antibody overnight at 4 °C. After washing with TBS-T, membranes were probed with horseradish peroxidase (HRP)-conjugated secondary antibodies. Signal detection was performed with the iBright FL1500 machine (Thermo Fisher Scientific, Waltham, MA) using Immobilon Super ECL (WBKLS0500, MilliporeSigma, St Louis, MO). Signal intensities were quantified by densitometry using ImageJ software (NIH, Bethesda, MD).

In other experiments, conditioned media was collected, concentrated using Amicon Ultra 2mL 50 kDa MWCO centrifugal filters (UFC205024, MilliporeSigma, St. Louis, MO) following the manufacturer’s instructions. Western blot analysis was then performed on concentrates after resuspension in Laemmli Buffer and boiling.

### IRF3 Immunoprecipitation and mass spectrometry

LX2 cells were cultured in 15-cm dishes and then stimulated with TGFβ for 16 hours. After washing with ice-cold PBS, cells were lysed in Purification buffer [50 mM Tris-HCl pH 7.4, 150 mM NaCl, 1 mM EDTA, 1% Triton X-100] supplemented with Protease and Phosphatase Inhibitors. Protein concentration was measured using DC Protein Assay. Co-immunoprecipitation was then performed with an equivalent amount of mouse anti-IRF3 antibody or mouse IgG1 isotype control. Lysate, antibody, and washed Pierce Protein A/G Magnetic Beads (88803, Thermo Fisher Scientific, Waltham, MA) were incubated in Purification buffer with rotation for four hours at 4 °C. After washing, immunoprecipitated proteins were eluted with 0.1 M glycine-HCl (pH 2.8). After neutralization of the acidic glycine with 1.5 M Tris (pH 8.8), immunoprecipitated proteins were resuspended in Laemmli Buffer + beta-mercaptoethanol, boiled at 95 °C for 5 minutes, then analyzed via Western blotting using light-chain specific secondary antibodies.

In the proteomics experiment in Figure 3, LX2 cells were cultured in 15-cm dishes. Unstimulated LX2 cells were lysed in Purification buffer, then immunoprecipitation was performed with mouse anti-IRF3 antibody. Immunoprecipitated beads were removed from magnetic beads by boiling in NuPage LDS Sample Buffer (NP0007, Thermo Fisher Scientific, Waltham, MA), then run on SDS-PAGE using 4-12% NuPage Bis-tris Mini Protein Gels (NP0321BOX, Thermo Fisher Scientific, Waltham, MA) using NuPAGE MOPS SDS Running Buffer (NP0001, Thermo Fisher Scientific, Waltham, MA). Silver staining was performed on the gel, and seven bands (Supplemental Figure 2) corresponding to areas with visible protein were cut out, digested, and subjected to mass spectrometry using Bruker TimsTof Pro2 Q-Tof mass spectrometry system. Gene Ontology analysis of identified proteins was then performed using Metascape^59^.

### Immunofluorescence

LX2 cells were plated in Lab-Tek Permanox Chamber Slide system (177445, Thermo Fisher Scientific, Waltham, MA) after pre-coating with poly-L-lysine (P4707, Sigma-Aldrich, St. Louis, MO). Poly(I:C) transfection or TGFβ stimulation was then performed. In certain experiments, cells were pre-treated for 24 hours with 2-CM (sc-283467, Santa Cruz, Dallas, TX). In certain experiments, cells were treated with MitoTracker Orange (M7510, Invitrogen, Waltham, MA) following the manufacturer’s instructions immediately prior to the end of the stimulation. Cells were then fixed with 4% paraformaldehyde, quenched with 25 mM glycine, permeabilized with 0.02% saponin, blocked with 10% goat serum in PBS for an hour at RT, then incubated with primary antibody overnight at 4 °C. After washing with PBS, cells were then incubated with secondary antibodies for an hour at RT, counterstained with DAPI (62248, Thermo Scientific, Waltham, MA), then mounted with Vectashield Vibrance Antifade Mounting Medium (H-1700, Vector Laboratories, Newark, CA). Z-stack images were acquired with 63x oil objective and a step size of 0.3 μm using a Leica TCS-SP8-AOBS inverted confocal microscope (Leica Microsystems, GmbH, Wetzlar, Germany). Z-stack images of 3-4 different fields (with each field containing approximately 15-30 cells) were obtained for each experimental condition.

### IMARIS analysis

Confocal Z-stacks were imported into the Imaris software (Bitplane, Belfast, United Kingdom). The *Surfaces* module, where a surface object is built around the structures of interest in a particular channel based on the fluorescent intensity, was used to define nuclei (using DAPI fluorescent channel), CAPRIN1^+^ or G3BP1^+^ stress granules (using anti-CAPRIN1 or anti-G3BP1 fluorescent channel), and mitochondria (using MitoTracker fluorescent channel). Next, the *Masked Channel* feature was used to define the anti-dsRNA (J2) or anti-IRF3 fluorescent signals inside and outside the previously-created surfaces (nuclei, stress granules, and mitochondria). Masked channels have values of zero in voxels lying outside of the surface and the value of the original channel within the surface (or vice versa). Multiple statistical parameters, including the volume and fluorescent intensity sums of each surface and both unmasked and masked channel were exported. For each surface or channel, the fluorescent intensity sums of each field were tabulated and normalized to the volume of nuclei within that field.

### Statistical analysis

Values in all figures represent mean ± standard error of the mean (SEM) unless otherwise indicated. Analysis of variance was performed using the general linear models procedure (SAS, Cary, NC). Data were log-transformed as necessary to obtain a normal distribution. Follow-up comparisons were made by least square means testing with p-values < 0.05 considered to be significant.

## DATA AVAILABILITY

All data generated or analyzed during this study are included in the main text or Supplementary Information. Source data are provided with this paper.

## ACKNOWLEDGEMENTS

Financial support: This work was supported in part by NIH grants: P50 AA024333 (to LEN and DMR), R01 AA027456 (to LEN), R01 NS112526 (to DD), K01 AA029474 (to JW), and F32 AA029290 (to CKD). This work was also supported by the Harrington Physician-Scientist Pathway of University Hospitals / Case Western Reserve University (to JT). The authors acknowledge the assistance of the Cleveland Clinic Lerner Research Institute Imaging Core in providing microscopy services. This work utilized the *Leica SP8* confocal microscope that was purchased with funding from National Institutes of Health SIG grant 1S10OD019972-01. The timsTof Pro2 instrument was purchased via an NIH shared instrument grant, S10 OD030398.

