## Supplementary Information for "Mitochondria-derived dsRNA-induced stress granules promote IRF3-mediated fibrotic responses"

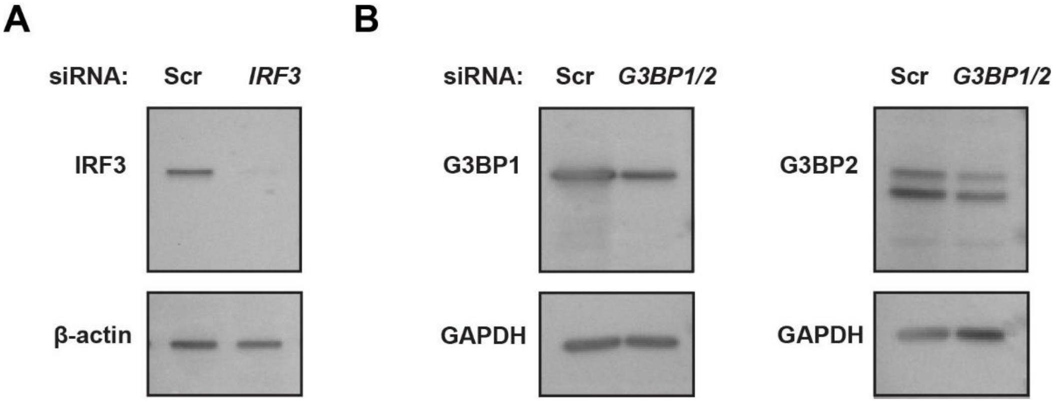


**Supplementary Figure 1: Confirmation of siRNA-mediated gene silencing in LX2 cells.** LX2 cells were transfected with scrambled control siRNA or siRNA against *IRF3* (**A**) or *G3BP1* and *G3BP2* (**B**). Western blot was performed on cell lysates to assess levels of indicated proteins. Images are representative of 1-2 independent experiments.

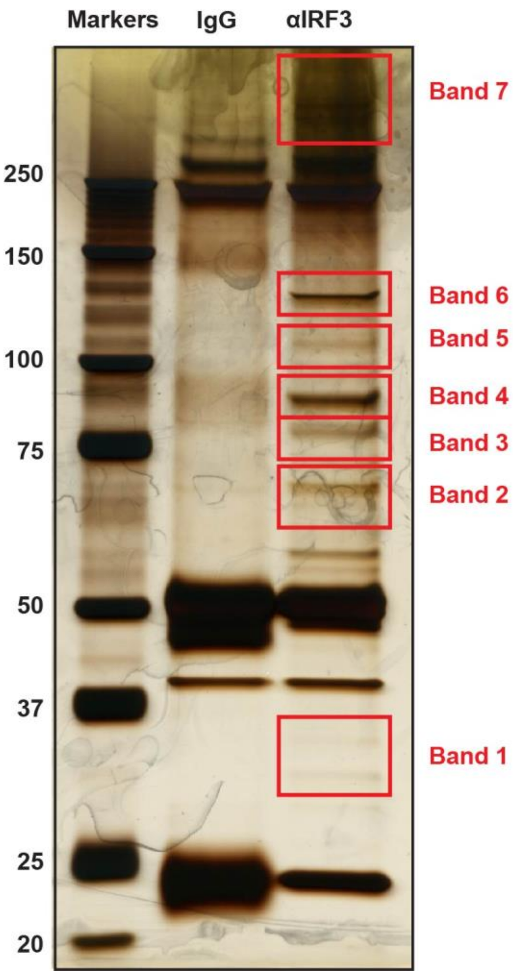


**Supplementary Figure 2: SDS-PAGE of immunoprecipitated proteins prior to mass spectrometry in LX2 cells.** Immunoprecipitation was performed with anti-IRF3 antibody in unstimulated LX2 cells, and immunoprecipitates were run on SDS-PAGE. The seven indicated bands were subjected to mass spectrometry.

**Supplementary Table 1: Summary of IRF3-binding proteins identified by mass spectrometry in LX2 cells.**

The identities of immunoprecipitated proteins identified by mass spectrometry are listed, as well as the number of total and unique peptides, protein sequence coverage percentage, and Spectral Abundance Factor (SAF).

**Supplementary Table 2: List of gene-specific primers used for quantitative real-time polymerase chain reaction.**
